# A gene silencing screen uncovers diverse tools for targeted gene repression in Arabidopsis

**DOI:** 10.1101/2022.11.01.514775

**Authors:** Ming Wang, Zhenhui Zhong, Javier Gallego-Bartolomé, Zheng Li, Suhua Feng, Peggy Hsuanyu Kuo, Ryan L. Kan, Hoiyan Lam, John Richey, Yasaman Jami-Alahmadi, James Wohlschlegel, Steven E. Jacobsen

**Author notes:** These authors contributed equally (Ming Wang, Zhenhui Zhong).

## Abstract

DNA methylation has been utilized for target gene silencing in plants, however it’s not well-understood whether other silencing pathways can be also used to manipulate gene expression. Here we performed a gain of function screen for proteins that could silence a target gene when fused to an artificial zinc finger. We uncovered many proteins that suppressed gene expression either through the establishment of DNA methylation, or via DNA methylation-independent processes including histone H3K27me3 deposition, H3K4me3 demethylation, H3K9, H3K14, H3K27, and H4K16 deacetylation, inhibition of RNA Polymerase II transcription elongation or Ser-5 dephosphorylation. The silencing fusion proteins also silenced many other genes with different efficacy, and a machine learning model could accurately predict the efficacy of each silencer based on various chromatin features of the target loci. These results provide a more comprehensive understanding of epigenetic regulatory pathways and provide an armament of tools for targeted manipulation of gene expression.

---

Transcriptional gene regulation is a fundamental biological process that controls the on or off states of gene expression, and involves DNA methylation, histone modification and chromatin remodeling ^1^. In plants, DNA methylation is generally linked to transcriptional gene silencing ^2^. For example, the Arabidopsis *FWA* gene is normally DNA methylated and silenced in all tissues, except in the developing endosperm where it is demethylated and expressed^3^.

In addition to DNA methylation, histone modifications also contribute to gene silencing ^1^. For example, Polycomb-Repressive Complex 1 (PRC1) and PRC2 are conserved in plants and animals and act to silence genes via the histone mark H3K27 trimethylation (me3) ^4–6^. Gene silencing can be also achieved by removing activating histone marks. For example, histone H3K4me3 and histone acetylation are associated with active gene activity, which can be erased through H3K4 demethylases such as JUMONJI containing proteins (JMJs), and histone deacetylases (HDACs), respectively ^7^, ^8^, ^9^.

Artificial zinc fingers are DNA binding domains that can be designed to bind a specific sequence and guide fusion proteins to specific loci ^10^. For example, artificial zinc finger 108 (here after ZF) was designed to bind the Arabidopsis *FWA* promoter in the region that is normally methylated in Col-0 wild type plants ^11^. When ZF was fused with the RNA-directed DNA methylation (RdDM) component SUVH9 and transformed into *fwa* epiallele containing plants, *FWA* DNA methylation and suppression were restored ^11^. It was later shown that ZF fusions with many other RdDM related proteins also caused *FWA* silencing and methylation ^12–14^. However, it’s largely unknown whether ZF fusions with non-DNA methylation related chromatin proteins can also trigger target gene silencing ^15^.

In this study, we fused a panel of 270 putative Arabidopsis chromatin proteins to ZF, and screened for fusions capable of silencing *FWA*. We identified 14 proteins capable of silencing through diverse mechanisms including establishment of DNA methylation, H3K27me3 deposition, H3K4me3 demethylation, H3K9, H3K14, H3K27 and H4K16 deacetylation, inhibition of Pol II transcriptional elongation, or Pol II dephosphorylation. We found that some target genes were only silenced by certain effector proteins, and a machine learning model could accurately predict which genes would be effectively silenced based on proximal chromatin features and expression levels of the target genes. These findings lay a comprehensive foundation for more detailed mechanistic understanding of gene silencing pathways and provide an array of new tools for targeted gene silencing.

## Results

### A gain of function screen for regulators of gene silencing in Arabidopsis

We utilized the native Arabidopsis gene *FWA* as a reporter to screen for regulators of gene silencing. *FWA* is a transcription factor gene that causes a late flowering phenotype when overexpressed, resulting in a greater number of leaves produced before flowering. In Col-0 wild type plants, *FWA* is completely silenced by DNA methylation, while in *fwa* epialleles which have permanently lost this DNA methylation *FWA* misexpression causes late flowering ^16^. In order to find putative gene silencing regulators, we searched the Arabidopsis ORFeome collections ^17, 18^ for chromatin related proteins, and also added other proteins of interest that were not present in the ORFeome collections (Supplementary Table 1). 270 putative Arabidopsis chromatin proteins were fused with a ZF designed to bind the *FWA* promoter ^11^. These fusions were individually transformed into *fwa* plants to screen for regulators that triggered *FWA* silencing and restored an early flowering phenotype.

This screen identified 14 effector proteins that successfully restored the early flowering phenotype of the *fwa* epiallele (Fig. 1a). Among these effectors, DMS3, SUVH2, SUVH9, and MORC1 are known players in the RdDM pathway, and previous studies have shown that DMS3-ZF, SUVH9-ZF, and MORC1-ZF could restore methylation and silencing of *FWA* ^11, 12^. SUVH2 is a close homolog of SUVH9 that functions in RdDM ^11^, and it was thus not unexpected that ZF-SUVH2 would also methylate and silence *FWA* expression (Fig. 1b-d).

**Fig. 1:**
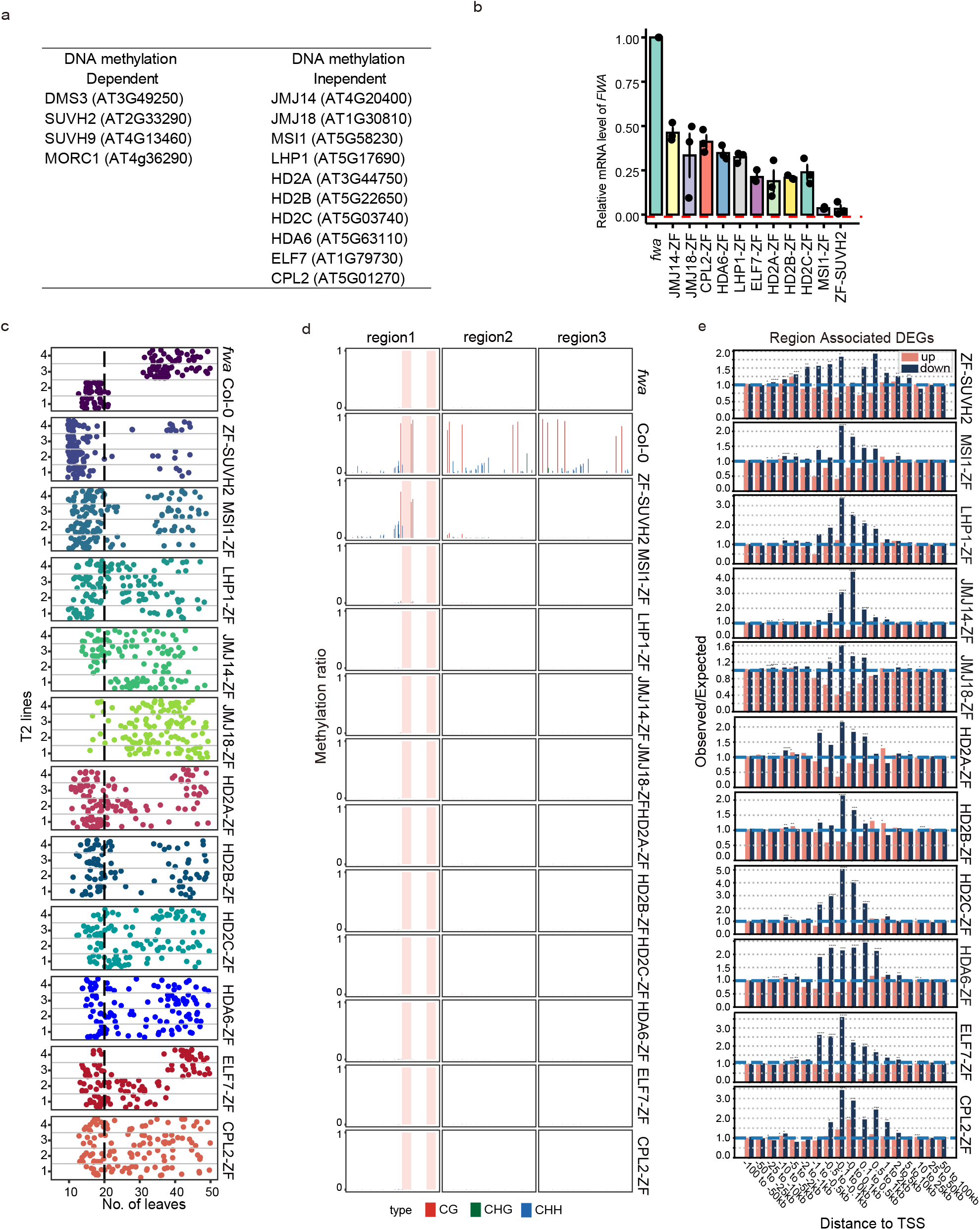
The effector proteins obtained from ZF target screening. **a,** The list of effector proteins identified from ZF target screening, which were dependent (left) or independent (right) of DNA methylation. **b,** Bar chart showing the relative mRNA level of *FWA* in *fwa* and three representative T2 ZF fusion lines using normalized reads of RNA-seq data (RPKM). **c,** Flowering time of *fwa*, Col-0 and four representative T2 ZF fusion lines as measured by the number of the leaves. **d,** CG, CHG, and CHH DNA methylation levels over *FWA* promoter regions in *fwa*, Col-0 and representative T2 ZF fusion lines measured by bisulfite (BS)-PCR-seq. Pink vertical boxes indicated ZF binding sites. **e,** The Observed/Expected values of up (pink bars)- and down (dark blue bars)-regulated DEGs over ZF off-target sites in ZF fusion lines, measured by Region Associated DEG (RAD) analysis. The asterisks indicated the *p* value calculated with hypergeometric test, *p < 0.05: \**; *p < 0.01: \*\**; *p < 0.001: \*\*\**; *p < 0.0001: \*\*\*\**.

We also identified many gene regulators that silenced *FWA* in a DNA methylation-independent manner (Fig. 1a-d), including two polycomb-group proteins MULTICOPY SUPRESSOR OF IRA1 (MSI1) and LIKE HETEROCHROMATIN PROTEIN 1 (LHP1), two JUMONJI domain containing proteins JMJ14 and JMJ18, four histone deacetylases HD2A, HD2B, HD2C, and HDA6, an RNA polymerase II-associated factor 1 (PAF1) homolog EARLY FLOWERING 7 (ELF7), and Carboxyl-terminal Domain Phosphatase-like 2 (CPL2) (Fig. 1a). The ZF fusions of these proteins restored an early flowering phenotype of the *fwa* epiallele to a similar level as in wild type Col-0 plants (Fig. 1c and Supplementary Fig. 1a), even though the *FWA* silencing was less efficient than ZF-SUVH2 (Fig. 1b and Supplementary Fig. 1b).

Although ZF was designed to bind the *FWA* promoter, it also binds to thousands of off-target sites throughout the genome ^12^. We therefore performed RNA-seq to determine if the different ZF fusion proteins could also regulate other genes near these off-target binding sites. We analyzed genes near 6,091 ZF ChIP-seq peaks that showed at least four-fold of enrichment of ZF ChIP-seq signal relative to a *fwa* non transgenic control (Supplementary Table 2). The differentially expressed genes (DEGs) near ZF off-target peaks were analyzed using Region Associated DEG (RAD) analysis ^19^. This analysis showed that all the ZF fusions showed a significantly higher number of downregulated DEGs than upregulated DEGs when the ZF peak was within one kilobase of the start site of the gene (Fig. 1e and Supplementary Fig. 1c), suggesting that all the identified fusions can repress many other genes in addition to *FWA*.

### Target gene silencing is non-heritable in DNA methylation independent ZF fusions

To test inheritability of the DNA methylation in ZF-SUVH2, bisulfite amplicon sequencing analysis (BS-PCR-seq) was performed to evaluate DNA methylation at *FWA* promoter regions in T2 lines that still contained the transgene and in lines that had segregated away the transgene (null segregants). DNA methylation was observed in both transgenic and null segregant lines, showing that DNA methylation established by ZF-SUVH2 was heritable in the absence of the transgene (Fig. 2a), as has been shown for other fusion proteins that target methylation to *FWA* ^11, 12, 20^. As expected, the early flowering phenotype was also inherited in many null segregant plants in the T2 population (Fig. 2b).

**Fig. 2:**
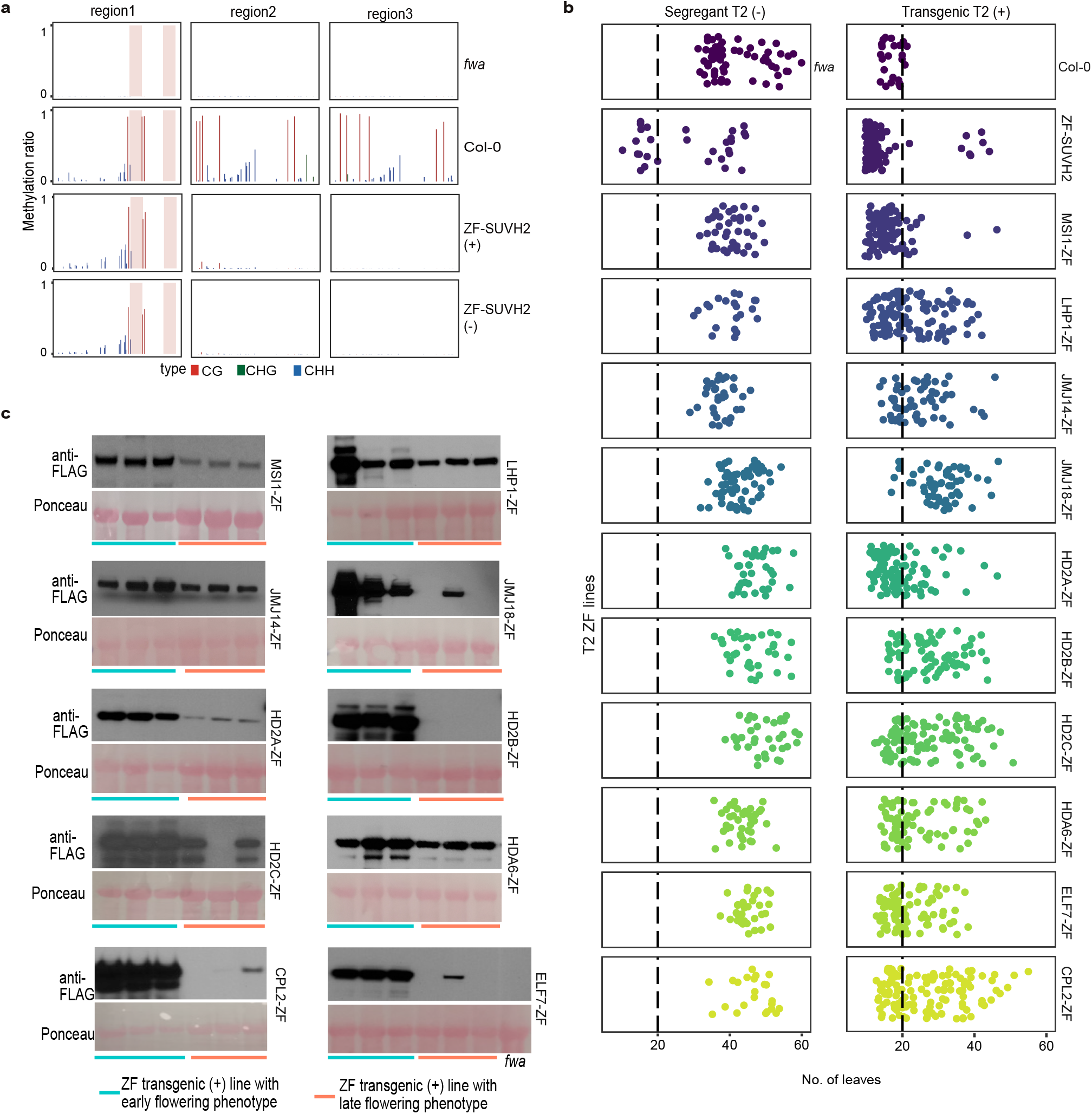
The inheritability of target gene silencing in the T2 lines of ZF fusions. **a,** CG, CHG, and CHH DNA methylation levels over *FWA* promoter regions in *fwa*, Col-0, ZF-SUVH2 transgenic T2 line (+), and ZF-SUVH2 segregant T2 line (-), measured by Bisulfite (BS)-PCR-seq. Pink vertical boxes indicated ZF binding sites. **b,** Flowering time of ZF fusions T2 segregant lines (-) and transgenic lines (+). **c,** Western blot showing the protein expression levels of ZF fusion T2 transgenic lines with early flowering phenotype (left three samples) and late flowering phenotype (right three samples).

We similarly analyzed the heritability of the flowering time phenotypes for the ZF fusions that were not associated with *FWA* DNA methylation. We found that in T2 plants that inherited the fusion protein transgenes, the early flowering phenotype was usually maintained (Fig. 2b). However, in all null segregant plants, the flowering time reverted to the typical late flowering phenotype of *fwa* plants (Fig. 2b), showing that the persistent presence of the fusion protein transgenes was needed for *FWA* silencing. Within the population of transgene containing T2 plants we observed wide variation in flowering time (Fig. 2b). This was likely due to differences in the expression level of the fusion proteins, as we observed by Western blotting that plants with high levels of transgene expression tended to have an early flowering phenotype, while plants with low protein expression levels tended to have a late flowering phenotype (Fig. 2c).

### Targeted gene silencing by LHP1, MSI1, and H3K27me3 deposition

MSI1 is a component of the PRC2 complex that also interacts with the Arabidopsis PRC1 component LHP1 ^21–23^, both of which are important for H3K27me3 mediated gene silencing. Therefore, to test whether H3K27me3 was deposited at *FWA* and ZF off-target sites, H3K27me3 and H3 ChIP-seq were performed in *fwa*, MSI1-ZF, and LHP1-ZF plants. Indeed, H3K27me3 ChIP-seq signals were higher at *FWA* in LHP1-ZF and MSI1-ZF than *fwa* control plants (Fig. 3a). We also observed H3K27me3 enrichment in LHP1-ZF and MSI1-ZF when plotting over 6,091 ZF off-target sites (Fig. 3b and Supplementary Fig. 2a), suggesting that tethering MSI1 and LHP1 to *FWA* and other target genes can cause gene silencing associated with H3K27me3 deposition.

**Fig. 3:**
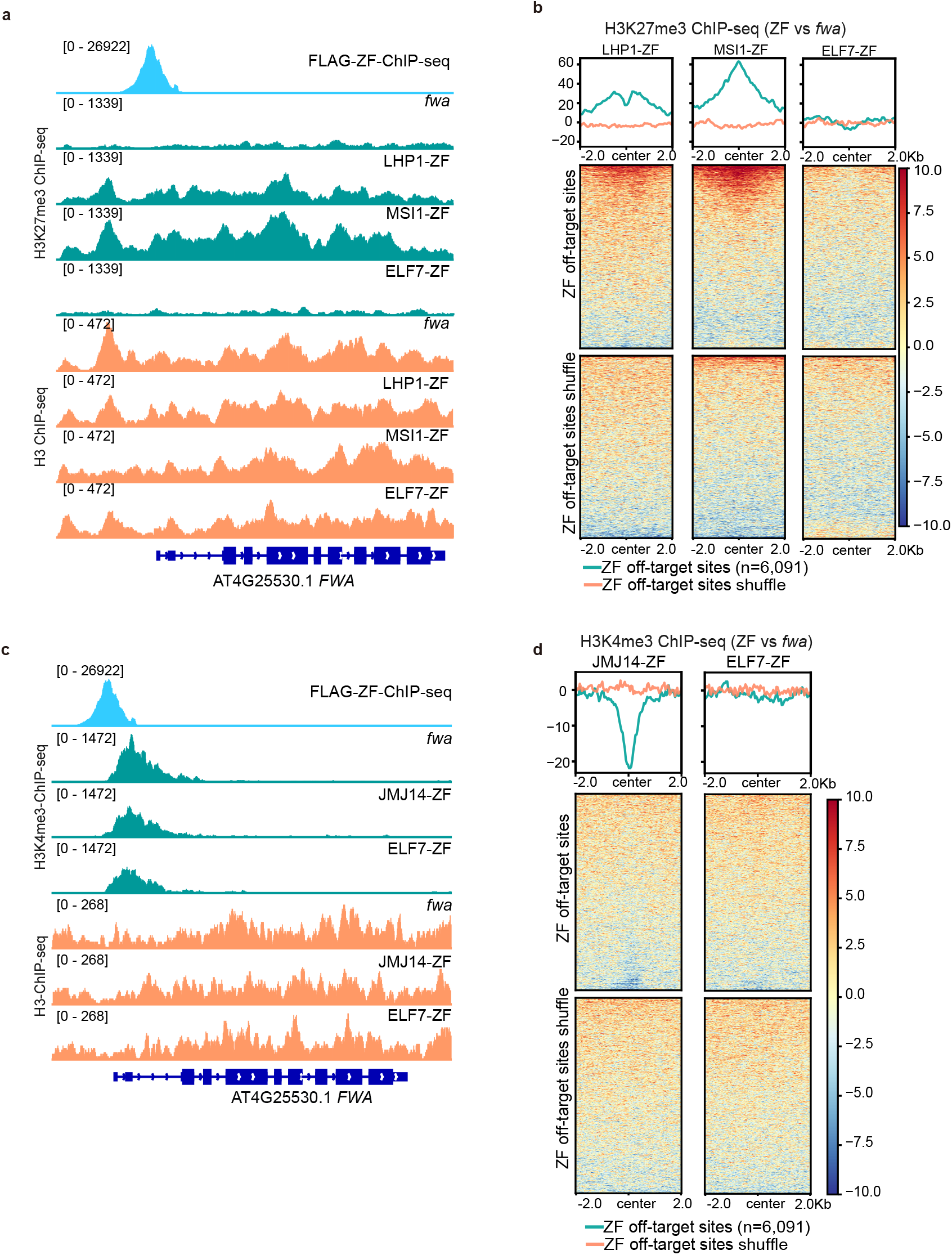
Targeted gene silencing by H3K27me3 deposition and H3K4me3 demethylation. **a,** Screenshots of H3K27me3 (up panel) and H3 (bottom panel) ChIP-seq signals over *FWA* region in *fwa*, LHP1-ZF, MSI1-ZF, and ELF7-ZF. FLAG-ZF ChIP-seq indicates ZF binding site. **b,** Metaplots and heatmaps depicting the normalized H3K27me3 ChIP-seq signals over ZF off-targeting sites (n=6,091) and ZF off-targeting sites shuffle in the representative T2 lines of LHP1-ZF, MSI1-ZF, and ELF7-ZF versus *fwa*, respectively. **c,** Screenshots of H3K4me3 (up panel) and H3 (bottom panel) ChIP-seq signals over *FWA* region in *fwa*, JMJ14-ZF and ELF7-ZF. The FLAG-ZF ChIP-seq signal indicates ZF binding site. **d,** Metaplots and heatmaps depicting the normalized H3K4me3 ChIP-seq signals in the representative T2 lines of JMJ14-ZF and ELF7-ZF over ZF off-targeting sites (n=6,091) and ZF off-targeting sites shuffle versus *fwa*, respectively.

### Targeted gene silencing by JMJ14, JMJ18, and H3K4me3 removal

Two H3K4 demethylase protein fusions, JMJ14-ZF and JMJ18-ZF, successfully triggered an early flowering phenotype and silenced *FWA* in a DNA methylation independent manner (Fig. 1a-e). H3K4me3 and H3 ChIP-seq was performed in JMJ14-ZF and an *fwa* control. We found that JMJ14-ZF caused a reduction of H3K4me3 over the *FWA* locus (Fig. 3c), as well over ZF off-target regions (Fig. 3d and Supplementary Fig. 2b). In addition, unlike MSI1-ZF and LHP1-ZF (Fig. 3a, b), JMJ14-ZF did not show accumulation of H3K27me3 at *FWA*, nor at ZF off-target regions (Supplementary Fig. 2c and d). Thus, silencing by JMJ14 is likely acting directly via removal of H3K4me3 rather than by accumulation of H3K27me3, a mark which can act antagonistically with H3K4me3 ^24–26^. Interestingly, several other H3K4 demethylase proteins were included in our collection including JMJ16/17, LDL1/2/3, and FLD, but none of these were able to trigger silencing of *FWA*.

### Targeted gene silencing by HDACs and histone deacetylation

Four histone deacetylase proteins, HD2A, HD2B, HD2C, and HDA6 were identified from the silencing screen. It was previously shown that Arabidopsis HD2A is required for H3K9 deacetylation and rRNA gene silencing ^27^, and that HD2C mediates H4K16 deacetylation and is involved in ribosome biogenesis ^28^, suggesting that HD2 family members can deacetylate multiple sites. We performed IP-MS utilizing a pHD2A:HD2A-FLAG transgene, which indicated that all three HD2 type HDACs interact with each other (Supplementary Table 3), consistent with an earlier report of IP-MS of tagged HD2C^28^.In addition, ChIP-seq analysis of pHD2A:HD2A-FLAG plants showed a partial overlap with histone H3K9ac, H3K27ac, and H4K16ac in the genome (Supplementary Fig. 3a, b). Thus, we profiled H3K9ac, H4K16ac, H3K27ac and H3 patterns by ChIP-seq in HD2A-ZF, HD2B-ZF, and HD2C-ZF (HD2-ZFs) plants. We found that H3K9ac, H4K16ac and H3K27ac were moderately reduced in HD2-ZF plants both at *FWA* (Fig. 4a) and over ZF off-target sites (Fig. 4b-d and Supplementary Fig. 4).

**Fig. 4:**
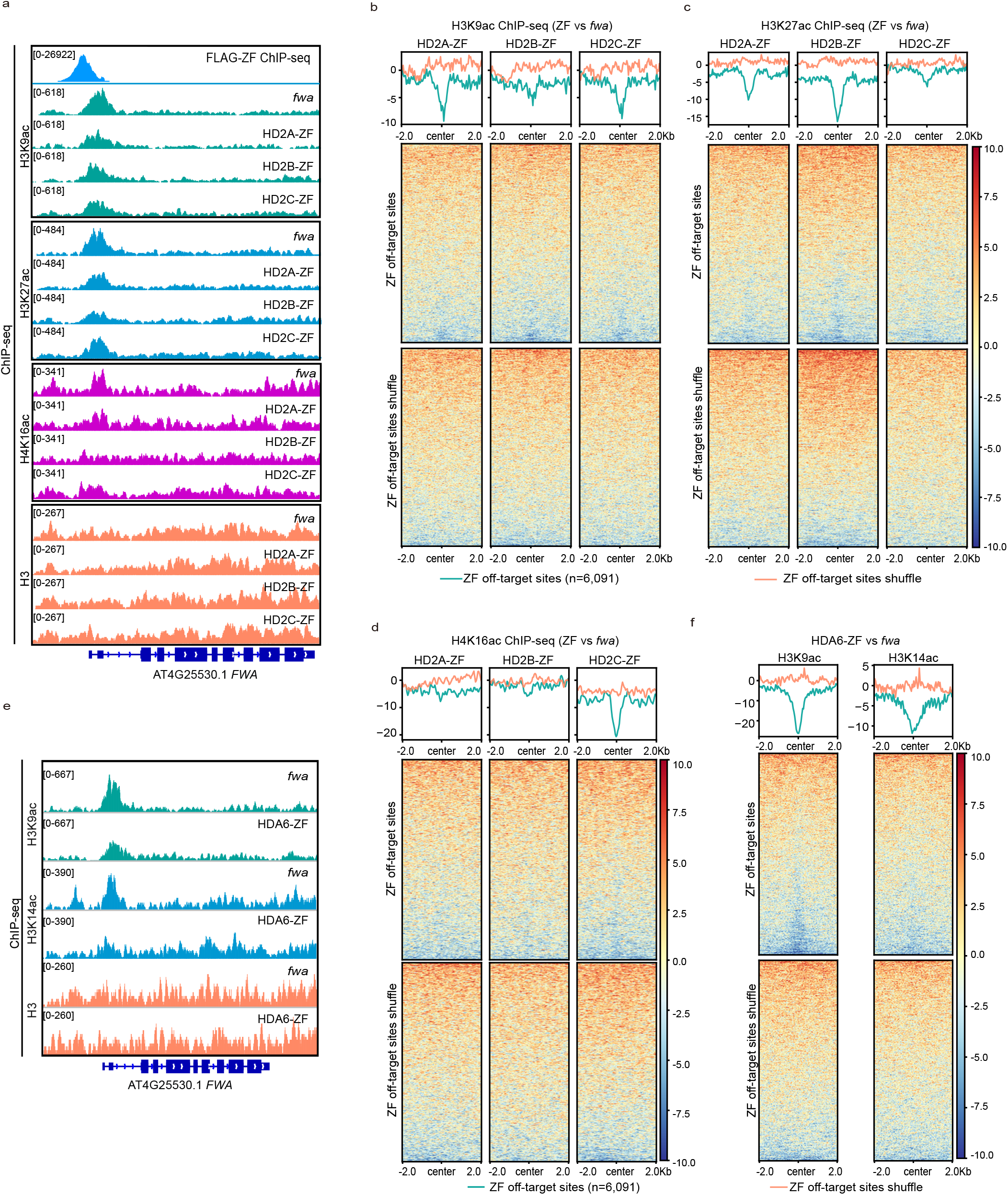
Targeted gene silencing by HDACs and histone deacetylation. **a,** Screenshots of histone H3K9ac, H3K27ac, H4K16ac, and H3 ChIP-seq signals over *FWA* region in *fwa*, HD2A-ZF, HD2B-ZF, and HD2C-ZF. **b-d,** Heatmaps and metaplots representing the normalized H3K9ac **(b)**, H3K27ac **(c)**, and H4K16ac **(d)** ChIP-seq signals over ZF off-target sites and ZF off-target sites shuffle in the representative T2 lines of HD2A-ZF, HD2B-ZF, and HD2C-ZF versus *fwa*. **e,** Screenshots of histone H3K9ac, H3K14ac, and H3 ChIP-seq signals over *FWA* region in *fwa* and HDA6-ZF. **f,** Heatmaps and metaplots representing the normalized H3K9ac and H3K14ac ChIP-seq signals over ZF off-target sites and ZF off-target sites shuffle in a HDA6-ZF T2 representative line versus *fwa*.

HDA6 has been reported to deacetylate several substrates including K9, K14, K18, K23, and K27 of the H3 histone tail and K5, K8, and K12 of the H4 histone tail ^29^, with H3K9ac and H3K14ac confirmed in multiple studies ^30, 31^. We therefore performed H3K9ac, H3K14ac, and H3 ChIP-seq in HDA6-ZF. Indeed, both H3K9ac and H3K14ac ChIP-seq signals were reduced at *FWA* as well as at ZF off-target sites in HDA6-ZF plants (Fig. 4e, f and Supplementary Fig. 5), suggesting that HDA6 represses target gene expression at least partially via histone H3K9 and H3K14 deacetylation. Together our results demonstrate that a variety of different histone deacetylase proteins of different classes can be harnessed for targeted gene silencing.

### Silencing by ELF7-ZF and interference with Pol II transcription

ELF7 encodes a PAF1 homolog, which is a subunit of the PAF1 complex (PAF1C). PAF1C is a conserved protein complex in eukaryotes that collaborates with RNA polymerase II during transcription initiation and elongation ^32, 33^. In Arabidopsis, mutation of PAF1C subunit *VIP3* caused a redistribution of histone H3K4me3 and H3K36me2 in certain genes ^34^, and we therefore initially performed H3K4me3, H3K36me2, and H3K36me3 ChIP-seq to see whether changes in these epigenetic marks might explain ELF7-ZF triggered *FWA* suppression. We observed some reduction of H3K4me3 at *FWA* in ELF7-ZF compared to *fwa* control plants (Fig. 3c). However, unlike JMJ14-ZF, H3K4me3 signal was largely unaffected near ZF off-target sites in ELF7-ZF (Fig. 3d). Considering that ELF7-ZF did trigger gene silencing at ZF off-target sites (Fig. 1e and Supplementary Fig. 1c), it seemed unlikely that H3K4me3 reduction was the relevant mechanism. In addition, signals of both H3K36me2 and H3K36me3 were slightly decreased at the *FWA* locus (Supplementary Fig.6a), while at the same time somewhat increased over ZF off-target sites (Supplementary Fig. 6b, c), making it unlikely that changes in H3K36me2 or H3K36me3 levels were the direct cause of ELF7-ZF mediated gene silencing.

We generated pELF7:ELF7-FLAG complementing transgenic lines in the *elf7-3* mutant background to perform IP-MS to identify ELF7 interacting proteins^35^ (Supplementary Fig. 7). Consistent with previous work ^33^ our ELF7 IP-MS data included peptides corresponding to all of the subunits of the PAF1C, as well as Pol II subunits and transcription factors, consistent with a role of ELF7 in Pol II transcription (Supplementary Table 3). Since ELF7 is a Pol II interacting protein, we hypothesized that ELF7-ZF might interact with Pol II at the *FWA* promoter region, retaining it there, and inhibiting transcription. To test this, we performed Pol II ChIP-seq in ELF7-ZF transgenic lines, as well as *fwa* and HD2A-ZF as controls. As expected, Pol II occupancy at *FWA* transcribed regions was significantly reduced in ELF7-ZF, as well as in HD2A-ZF (Fig. 5a), consistent with the silencing of *FWA* expression in these lines (Fig. 1b). However, we observed a very prominent Pol II peak at the *FWA* promoter overlapping the ZF site in the ELF7-ZF line, but not in HD2A-ZF nor *fwa* plants (Fig. 5a). Moreover, strong Pol II enrichment was also observed at the ZF off-target binding sites in ELF7-ZF (Fig. 5b and Supplementary Fig. 8a). Thus, Pol II appears to be tethered to the ZF binding sites via interaction with ELF7-ZF, which in turn appears to inhibit Pol II transcription, leading to gene silencing.

**Fig. 5:**
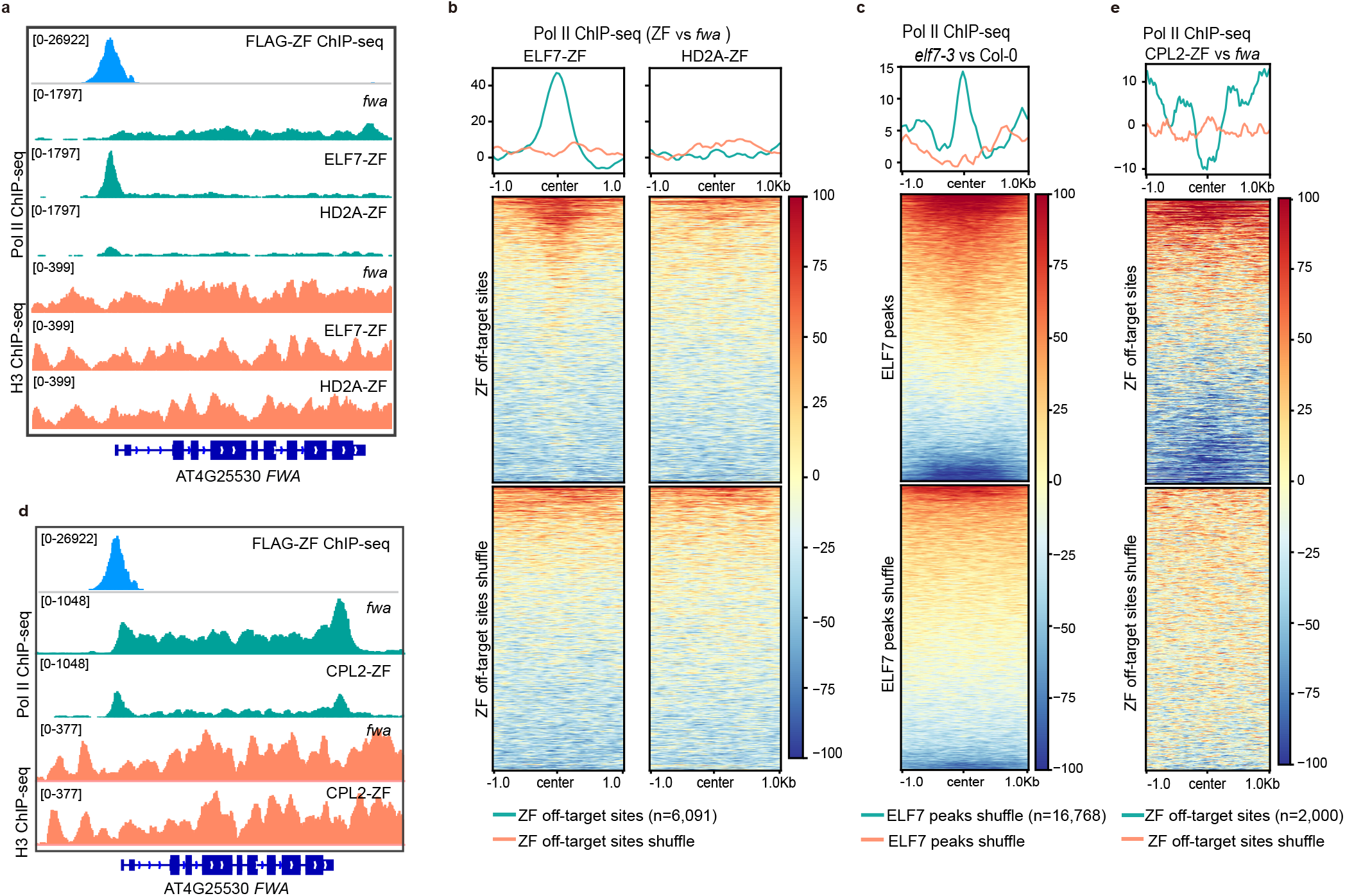
Targeted gene silencing by ELF7, CPL2, and Pol II transcription disruption. **a,** Screenshots of Pol II and H3 ChIP-seq signals over *FWA* region in *fwa*, ELF7-ZF, and HD2A-ZF. **b,** Heatmaps and metaplots representing normalized Pol II ChIP-seq signals in the representative T2 lines of ELF7-ZF and HD2A-ZF versus *fwa*, over ZF off-target sites (n=6,091) and ZF off-target sites shuffle, respectively. **c,** Heatmap and metaplot showing normalized Pol II ChIP-seq signals over ELF7-FLAG ChIP-seq peaks (n=16,768) and ELF7 peaks shuffle in *elf7-3* versus Col-0. **d,** Screenshots of Pol II ChIP-seq signals over *FWA* in *fwa* and a CPL2-ZF representative T2 transgenic line. **e,** Heatmap and metaplot depicting the normalized Pol II ChIP-seq signals over ZF off-target sites that had preexisting Pol II Ser5 ChIP-seq signals (n=2,000) and shuffle sites in CPL2-ZF versus *fwa*.

To better understand the endogenous function of ELF7, we performed ChIP-seq in pELF7:ELF7-FLAG transgenic lines. Consistent with its role in transcriptional elongation, ELF7 was exclusively distributed over gene body regions, with most ELF7 signals overlapping with both Pol II peaks and H3K36me2 or H3K36me3 peaks (Supplementary Fig. 8b, c). We also performed Pol II ChIP-seq in *elf7-3* and found a significant accumulation of Pol II at ELF7 enriched sites (Fig. 5c and Supplementary Fig. 8d), suggesting that transcriptional elongation is impeded, resulting in a higher Pol II occupancy reflected in the ChIP-seq data. Together our data show that the Arabidopsis PAF1C is required for proper Pol II transcriptional elongation, as has been shown in yeast and animal systems ^36, 37^, and that tethering the ELF7 component of the complex to promoters represents a novel synthetic mechanism to induce gene silencing that is likely independent of changes of particular epigenetic marks.

### Target silencing by CPL2-ZF and Pol II CTD Ser5 dephosphorylation

CPL2 is a well characterized phosphatase that specifically acts on serine 5 (Ser5) of the Pol II C-terminal domain ^38^, and represses transcription through inhibiting Pol II activity ^39, 40^. We therefore performed Pol II Ser5 ChIP-seq in CPL2-ZF transgenic lines, and we indeed observed reduced signal at *FWA* as well as ZF off-target sites that had preexisting Pol II Ser5 (Fig. 5d-e and Supplementary Fig. 9), suggesting that CPL2-ZF indeed silenced target genes through Pol II CTD Ser5 dephosphorylation. The promoter tethering of CPL2 thus represents a new mechanism for targeted gene silencing.

### Target genes vary widely in their sensitivity to different gene silencing approaches

We observed that the set of downregulated genes at ZF off-target sites for each ZF-fusion were partially non-overlapping (Supplementary Fig.10). As one example, gene AT3G13470 was downregulated by HD2A-ZF, HD2B-ZF, HD2C-ZF, and LHP1-ZF, but not ELF7-ZF nor CPL2-ZF (Fig. 6a). These results suggest that the best gene silencing approach will greatly depend on the particular target gene of interest, highlighting the utility of gene silencing tools that work by different mechanisms. We hypothesized that preexisting epigenetic features of the target genes might determine their sensitivity to silencing by the different ZF fusion proteins, and indeed different target genes showed different levels of various epigenetic marks including histone acetylation (H3K9ac, H3K27ac, and H4K16ac), histone methylation (H3K4me3 and H3K27me3), chromatin accessibility (ATAC-seq), and DNA methylation (Supplementary Fig. 11 and Supplementary Table 4). To test this hypothesis, we used the chromatin features of 2,709 genes containing nearby ZF binding sites, as well as the expression level of these genes, together with the information of whether or not these genes were silenced with each of the ZF-fusion as inputs to various machine learning algorithms (Fig. 6b and Supplementary Table 4). The K Neighbors Classifier showed the best performance and a 10-fold cross validation of this model showed an extremely high accuracy (82-98%) in predicting the efficacy of each silencer (Fig. 6b-c). This analysis supports the hypothesis that different effector proteins are efficient in silencing genes with different chromatin features.

**Fig. 6:**
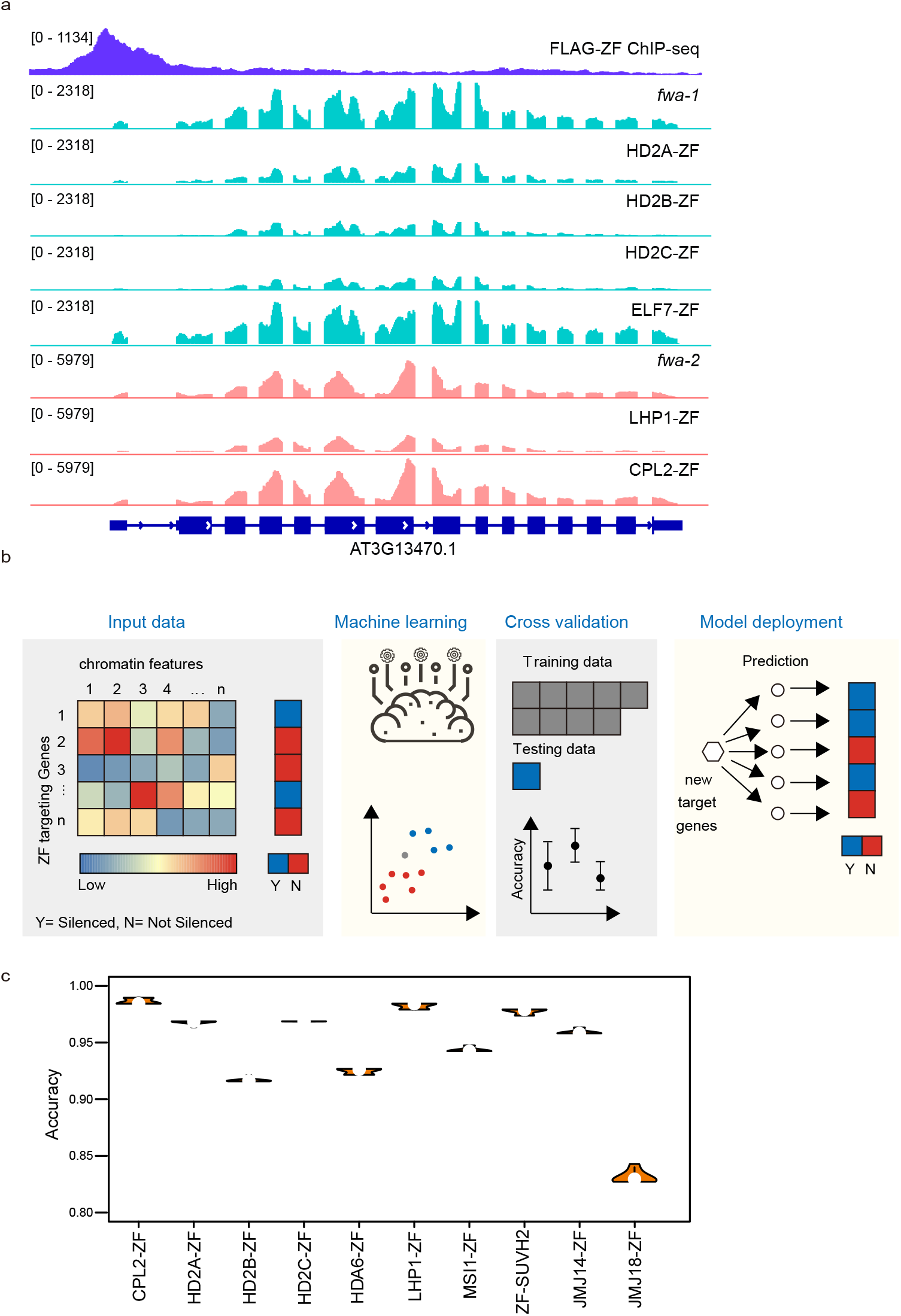
Predict the efficacy of different effector proteins by using machine learning models. **a,** Screenshots displaying RNA-seq levels of a representative ZF off-target gene AT3G13470 in *fwa-1*, HD2A-ZF, HD2B-ZF, HD2C-ZF, ELF7-ZF, *fwa-2*, LHP1-ZF and CPL2-ZF. FLAG-ZF ChIP-seq indicates the binding site of ZF. **b,** Flow charts showing the method of machine learning used to generate models for effector protein efficacy predictions. Y signifies that the ZF target genes were silenced in ZF lines, while N signifies that the ZF target genes were not silenced in ZF lines. **c,** Violin plot indicating the accuracy of cross validation in each ZF lines.

## Discussion

By constructing ZF fusions to target a collection of putative chromatin regulators to *FWA*, we uncovered a variety of proteins capable of inducing gene silencing at *FWA* as well as at many other loci. Our genomic and genetic evidence demonstrate that target gene silencing can be directed by diverse histone modifications including H3K27me3 deposition, H3K4me3 demethylation, and histone acetylation at H3K9, H3K14, and H4K16 tails. Interestingly, we also found two factors that appear to act by more directly interacting with Pol II, the ELF7 component of the elongation factor PAF1C and the CPL2 enzyme which dephosphorylates the RNA Pol II C-terminus.

Our results show that diverse pathways can be harnessed for the development of synthetic biology tools to downregulate genes. DNA methylation represents a strong and potentially heritable type of silencing, but only some genes will be amenable to this type of modification due to low densities of CG dinucleotides that are needed for silencing and heritability ^12^ or high levels of endogenous expression that can compete with methylation maintenance ^14^. In addition, it may often be desirable to cause only a partial silencing of a target gene, and the non-DNA methylation-based tools described here were able to cause various levels of repression that were lower than that of DNA methylation targeting tools (Fig.1b).

Our data also showed that certain genes were more amenable to particular gene silencing approaches, meaning that having a wider array of silencing tools expands the range of genes that can be successfully targeted. The genes susceptible to silencing in this study were involved in a wide range of biological processes including development, hormone signaling pathways, and disease resistance (Supplementary Table 4), suggesting that the tools described here could be useful for the modulation of many different traits. In addition, we were able to utilize machine learning algorithms which could accurately predict which genes would be sensitive to silencing by a particular gene silencing mechanism, which should be useful for future gene expression engineering efforts. In conclusion, this work provides mechanistic detail for an array of key plant gene silencing pathways, and describes a collection of new tools that should be useful in both basic research and crop improvement.

## Methods

### Plant materials and growth conditions

All the plants used in this paper were in the Arabidopsis thaliana Col-0 ecotype, grown under long-day conditions (16h light and 8h dark). The T-DNA insertion lines used in this study included *elf7-3* (SALK_019433). All the transgenic plants were generated by Agrobacterium (AGL0 strain) mediated floral dipping.

### Plasmid construction

**pMDC123-UBQ10: ZF-3XFLAG-EFFECTOR (cDNA)** is a Gateway compatible binary destination vector that includes a plant UBQ10 promoter, followed by an N-terminal ZF and 3XFLAG epitope tag, a Gateway cassette, and an OCS terminator. The list of selected effectors from Arabidopsis Gene ORFeome Collection (Supplementary Table 1) in the pENTR/D-TOPO vectors were all cloned into pMDC123 destination vector via LR reaction using Gateway LR Clonase II (Invitrogen). **pEG302-EFFECTOR (gDNA)-3XFLAG-ZF** is also a Gateway-compatible binary destination vector, which consists of a gateway cassette, followed by a C-terminal 3XFLAG epitope tag, a ZF, a Biotin Ligase Recognition Peptide (BLRP), and an OCS terminator. The sequence of native promoter (~1.5kb upstream from the 5’UTR or until the next gene annotation) and genomic DNA (No stop codon) of the effectors were cloned into pENTR D-TOPO vectors (Invitrogen), which were used to deliver the genomic DNA sequences of these effectors into the destination vector using Gateway LR Clonase II (Invitrogen). **pEG302-EFFECTOR (gDNA)-3XFLAG/9XMyc**: contains a gateway cassette, followed by a C-terminal 3XFLAG or 9xMyc epitope tag, a BLRP, and an OCS terminator. The cloning method is the same as pEG302-EFFECTOR (gDNA)-3XFLAG-ZF. Please see Supplementary Table 5 for the primers.

### Flowering time measurement

The flowering times were measured by the leaf counts, and each dot in the dot plots represented the leaf number of individual plants. We consider the plants with 20 or less leaves as early flowering.

### BS-PCR-seq

The leaf tissue from 4- to 5-week-old Col-0 wild type, *fwa*, and the representative T2 ZF lines showing early flowering phenotype were collected to perform Bisulfite PCR at *FWA* promoter regions. CTAB-based method was used to extract DNA and the EpiTect Bisulfite kit (QIAGEN) was used for DNA conversion. The converted DNA was used as a template to amplify three different regions over promoter and 5’ transcribed regions of *FWA*, including Region 1 (chr4: 13038143-13038272), Region 2 (chr4: 13038356- 13038499) and Region3 (chr4: 13038568-13038695). Pfu Turbo Cx (Agilent), dNTP (Takara Bio), and the primers designed for the above-mentioned *FWA* regions (see Supplementary Table 5) were used to perform PCR reactions. Three different PCR products from three regions of each sample were pooled and purified with AMPure beads (Beckman Coulter). The purified PCR products were used to construct libraries by the Kapa DNA Hyper Kit (Roche) together with TruSeq DNA UD indexes for Illumina (Illumina), and the libraries were sequenced on Illumina iSeq 100.

### IP-MS

The method of IP-MS used in this study has been described in a recent paper ^13^. Ten grams of unopened floral buds from Col-0 wild type and FLAG-tag transgenic plants, HD2A, and ELF7 were collected and ground into fine powder with liquid nitrogen. These samples were resuspended with 25mL IP buffer and homogenized until lump-free by dounce homogenizer. The lysate was filtered through Miracloth and incubated with 250μL anti-FLAG M2 magnetic beads (Sigma) at 4°C for 2 hours. The magnetic beads were washed with IP buffer and eluted with TBS containing 250 μg/mL 3X FLAG peptides. The eluted proteins were precipitated with trichloroacetic acid (Sigma) and subject to MS analyses as described previously ^13^.

### ChIP-seq

We followed previous protocol for ChIP-seq with minor modifications ^11^. Briefly, a total of 2-4 grams of leaves or unopened flower buds were collected. The plant materials were ground with liquid nitrogen and fixed with 1% formaldehyde containing nuclei isolation buffering for 10 minutes before adding fresh-made glycine to terminate the crosslinking reaction. The nuclei were isolated and disrupted by SDS, and the chromatin was sheared via Bioruptor Plus (Diagenode) and immunoprecipitated with antibody at 4°C overnight. Next, the magnetic Protein A and Protein G Dynabeads (Invitrogen) were added and incubated at 4°C for 2 hours. After washing and elution, the reverse crosslinking was done at 65°C overnight. Then the protein-DNA complex was treated with Protease K (Invitrogen) at 45°C for 4 hours, and the DNA was purified and precipitated with 3M Sodium Acetate (Invitrogen), GlycoBlue (Invitrogen) and Ethanol at −20°C overnight. The precipitated DNA was directly used for library construction using the Ovation Ultra Low System V2 kit (NuGEN), and the libraries were sequenced on Illumina NovaSeq 6000 or HiSeq 4000 instruments.

### RNA-seq

One leaf tissue of 4- to 5-week-old plants with similar age from *fwa* and early flowering ZF transgenic lines were collected for RNA extraction using Direct-zol RNA MiniPrep kit (Zymo Research). 1μg of total RNA was used to prepare the libraries for RNA-seq following TruSeq Stranded mRNA kit (Illumina), and the libraries were sequenced on Illumina NovaSeq 6000 or HiSeq 4000 instruments.

## Bioinformatic analysis

### BS-PCR-seq analysis

BS-PCR-seq data analysis in this study used the pipeline described in ^9^. The raw pair-end sequencing reads of each sample were combined, and aligned to both strands of reference genome TAIR10 using BSMAP (v.2.90) ^41^, and the alignment allowed up to 2 mismatches and 1 best hit. The reads with less than 20 reads coverage of cytosines and the reads with more than 3 consecutives methylated CHH sites were removed. The methylation level of each cytosine was calculated using the ratio of C/(C+T), and only the methylation data within the designed *FWA* regions was kept making a plot using customized R scripts.

### ChIP-seq analysis

The ChIP-seq raw reads were trimmed using trim_galore (https://www.bioinformatics.babraham.ac.uk/projects/trim_galore/) and then aligned to TAIR10 genome using bowtie version 1.1.2 ^42^, which allowed one unique mapping site 0 mismatch. The Samtools version 1.9 ^43^ was used to remove the duplicated reads, and together with deeptools version 3.1.3 ^44^ to generate tracks using RPKM for the normalization. The peaks were called using MACS2 version 2.1.1. ^45^, and the peaks that were frequently existed in previous FLAG ChIP-seq of Col-0 were removed.

For FLAG-ZF ChIP-seq, the FLAG ChIP-seq was performed in the unopened flower buds of FLAG-ZF T2 transgenic plants and *fwa* plants. The peaks were called by FLAG-ZF against *fwa*, and the peaks with 4 folds or higher signal enrichment were kept as the ZF off-target sites (Supplementary Table 2, n=6,091), while the other FLAG ChIP-seq used signal enrichment of 2 folds or higher for the following analysis.

For the comparison of Histone and Pol II (Histone/Pol II) enrichment over ZF off-target sites between ZF lines and *fwa*, the Histone/Pol II ChIP-seq of each sample including both ZF lines and *fwa* were normalized with their respective H3 ChIP-seq first by using bigwigCompare, and then the normalized Histone/Pol II ChIP-seq of ZF lines were further normalized to *fwa* by using bigwigCompare, which were then used to make the metaplot over ZF off-target peaks and random shuffle peaks. This method was also applied in Pol II ChIP-seq enrichments between Col-0 wild type and *elf7-3* mutant over ELF7 peaks and shuffle (Fig. 5c).

### RNA-seq analysis

The RNA-seq raw reads were aligned to TAIR10 genome using bowtie2 ^46^ and the expression levels were calculated with rsem-calculate-expression from RSEM with default settings ^47^. The RNA-seq tracks were generated using Samtools version 1.9 ^43^ and normalized with RPKM using bamCoverage from deeptools version 3.1.3 ^44^. The DEGs were called using customized scripts of run_DE_analysis.pl from Trinity version 2.8.5 ^48^. log2 FC ≥ 1 and FDR < 0.05 were used as a cut off.

The method of region associated DEGs analysis in Fig. 1e was described at ^14^. We used the FLAG-ZF ChIP-seq peaks (n=6,091) as input of the favorite regions, and up- and down-regulated DEGs of ZF lines versus *fwa* were used respectively as the inputs of DEGs.

### Machine learning

ZF targeting genes: We defined 6,091 ZF off target sites above, however, not all of them were located proximal to genes. Therefore, we considered 2,709 genes as ZF targeting genes, whose transcription start sites (TSSs) were located between −500bp to 200 from a ZF peak. The chromatin features of these 2,709 ZF target sites as well as the gene expression level of ZF target genes were subjected to the following analysis.

Input data preparation: A total of 15 chromatin or genomic features of the 2,709 ZF off target sites/genes, including gene expression level, ChIP-seq signals of H3K27ac, H3K27me3, H3K4me3, H4K16ac, H3K9ac, and Pol II, ATAC-seq signals, the level of CG, CHG, and CHH methylation, the number of CG, CHG, and CHH sites, and GC content, that have been characterized in the *fwa* epiallele (Supplementary Table 4) were utilized to generate input data for machine learning. We also used a binary gene classification based on whether the gene was significantly down-regulated (Fold Change < =2 and FDR < 0.05) in ZF lines compared with the *fwa* epiallele plants. Input data were preprocessed by removing perfect collinearity.

Prediction model selection: We evaluated 14 machine learning algorithms with 10-fold cross variation (Fig. 6c) in PyCaret (v2.3.4, https://pycaret.org/) to select prediction models. The input dataset was divided into 10 groups, with 9 groups used as training dataset and 1 group as test data, reiterated 10 times. The K Neighbors Classifier showed the best performance and was therefore applied in modeling in 13 ZF fusion protein lines. Hyperparameters of the created K Neighbors Classifier were further tuned to improve accuracy.

## Data availability

The accession number for all the high-throughput sequencing data of this paper is GEO: GSE197XXX.

## Acknowledgements

We thank Soo-Young Park, Benjamin Feng and Jiachen Wu for technical support; Dr. Colette Picard, Dr. Wanlu Liu and Dr. Ranjith Papareddy for bioinformatics advice. We also thank Mahnaz Akhavan and the UCLA BSCRC BioSequencing Core for the sequencing support. This work was supported by a NIH grant (R35GM130272) and a Bill and Melinda Gates Foundation grant (OPP1125410) to S.E.J. S.E.J. is Howard Hughes Medical Institute Investigator.

## Author Contributions

M.W. and S.E.J. designed the research, interpret the data, and wrote the manuscript; M.W. performed most of the experiments with the help from J.G.B., Z.L., P.H.K., R.K., H.Y.L., and J.R. Z.Z and MW performed bioinformatic data analysis; M.W., Y.J.A., and J.W performed IP-MS and interpreted the data. S.F. performed BS-PCR-seq and high throughput sequencing.

