## Supplement for "A gene silencing screen uncovers diverse tools for targeted gene repression in Arabidopsis"

Supplementary Fig. 1-11

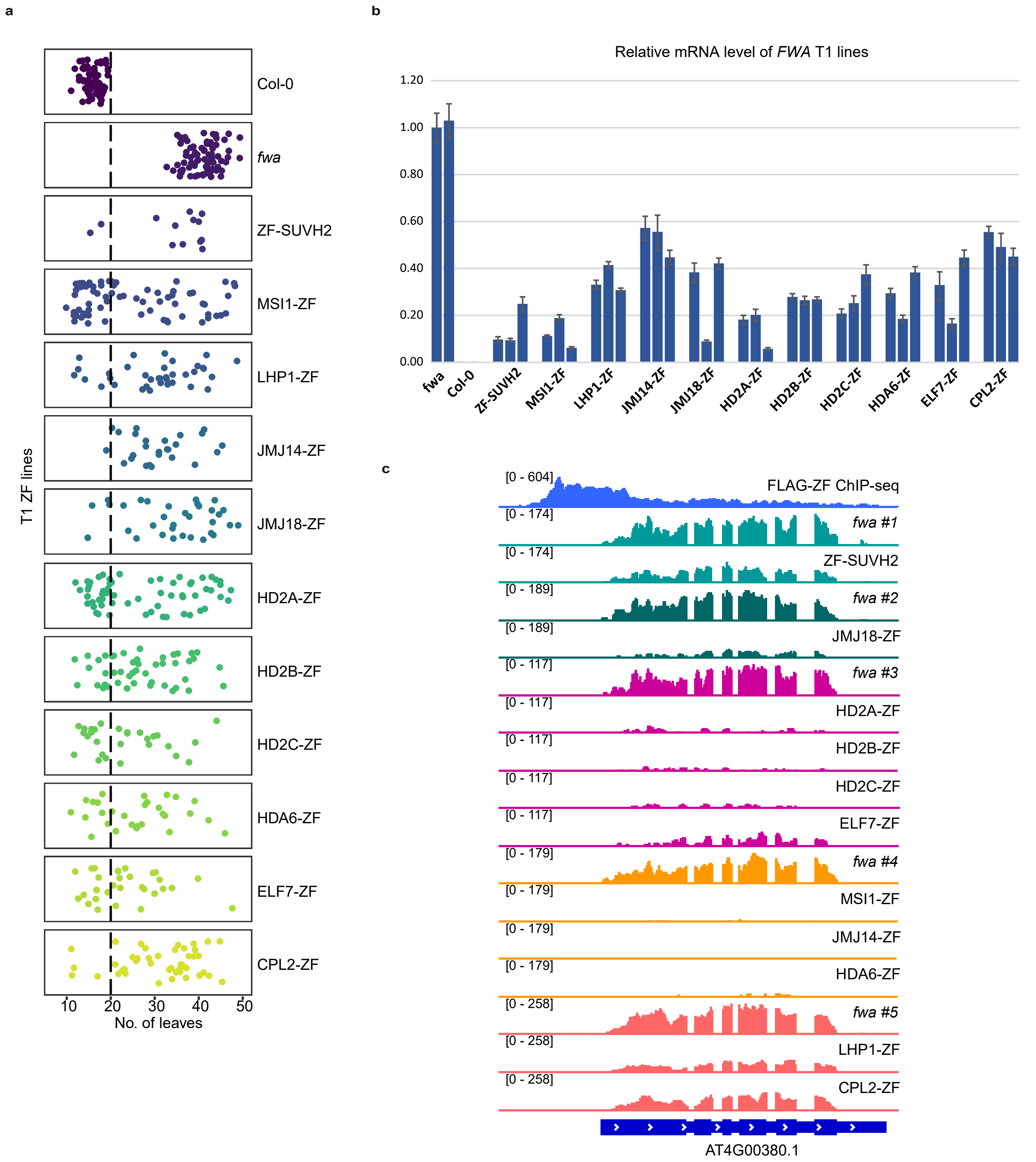


**Supplementary Fig. 1: ZF screening identified silencers. a,** Flowering time of *fwa*, Col-0, and T1 lines of ZF fusions. **b,** qRT-PCR showing the relative mRNA level of *FWA* gene in *fwa*, Col-0 and three representative T1 lines of ZF fusions. Error bars indicate standard error of three technical replicates. **c,** Screenshots of RNA-seq signals in *fwa* and representative T2 transgenic lines of ZF fusions over a representative ZF off-target site. The FLAG-ZF ChIP-seq indicates ZF binding site.

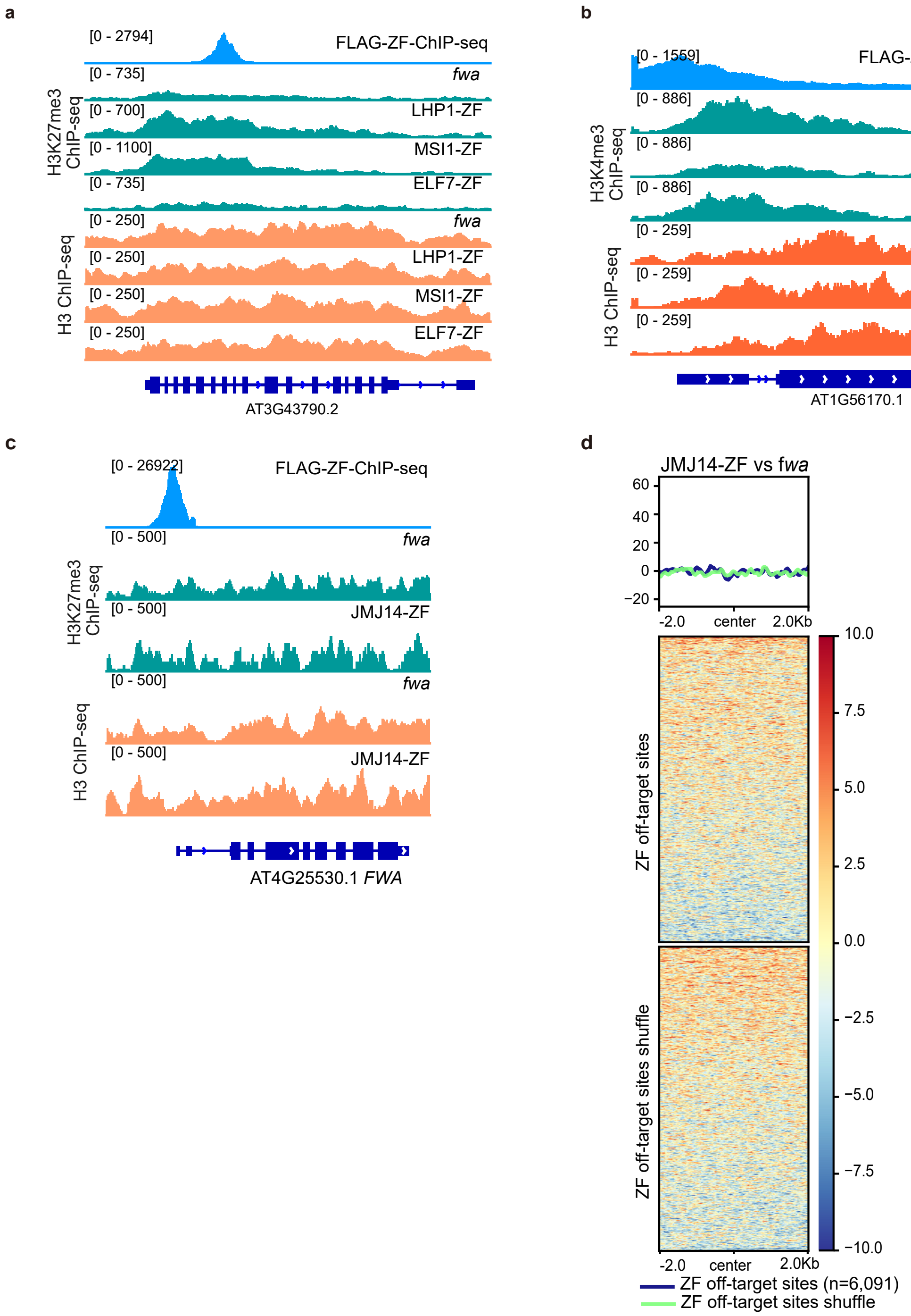


**Supplementary Fig. 2: Target gene silencing by histone H3K27me3 deposition and H3K4me3 demethylation. a,** Screenshots of H3K27me3 and H3 ChIP-seq signals over a representative ZF off-target site in *fwa* and T2 transgenic lines of LHP1-ZF, MSI1-ZF, and ELF7-ZF. **b,** Screenshots of H3K4me3 and H3 ChIP-seq signals in *fwa* and T2 transgenic lines of JMJ14-ZF and ELF7-ZF over a representative ZF off-target site. The FLAG-ZF ChIP-seq signals indicate ZF binding sites. **c,** Screenshots of H3K27me3 and H3 ChIP-seq signals in *fwa* and a representative T2 line of JMJ14-ZF over *FWA*. **d,** Heatmaps and metaplot showing the normalized H3K27me3 ChIP-seq signals over ZF off-target sites (n=6,091) in JMJ14-ZF T2 transgenic line versus *fwa*.

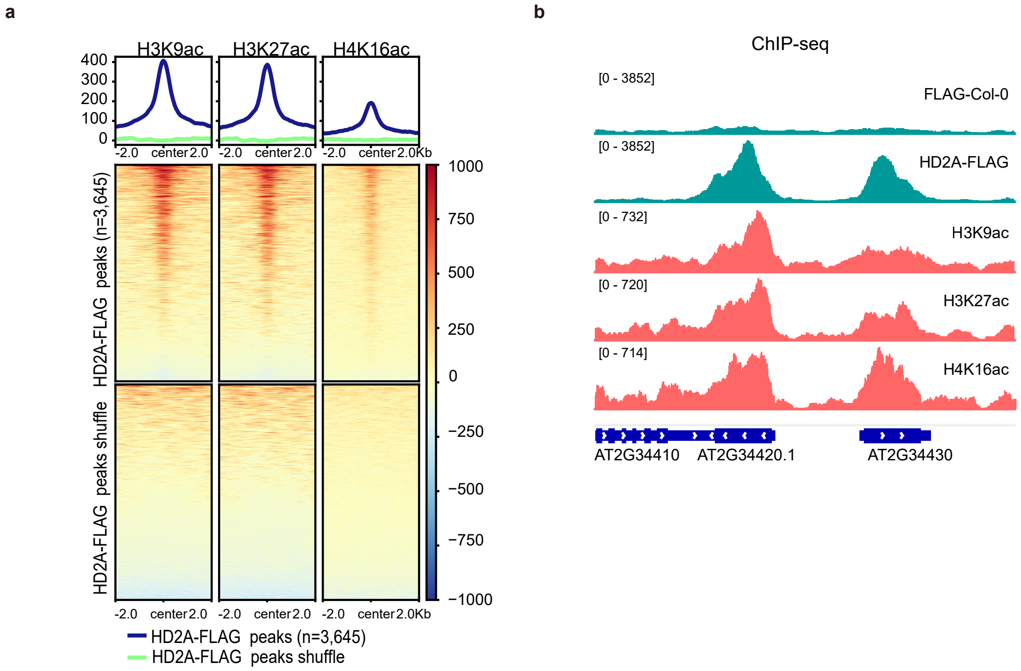


**Supplementary Fig. 3: HD2A overlapped with histone H3K9ac, H3K27ac, and H4K16ac. a,** Heatmaps and metaplots representing the H3K9ac, H3K27ac, and H4K16ac ChIP-seq signals over HD2A-FLAG ChIP-seq peaks (n=3,645, top panel) and shuffle sites (bottom panel). **b,** Screenshots showing the FLAG ChIP-seq signals in Col-0, HD2A-FLAG, and histone H3K9ac, H3K27ac, and H4K16ac ChIP-seq signals in Col-0 over two representative genes.

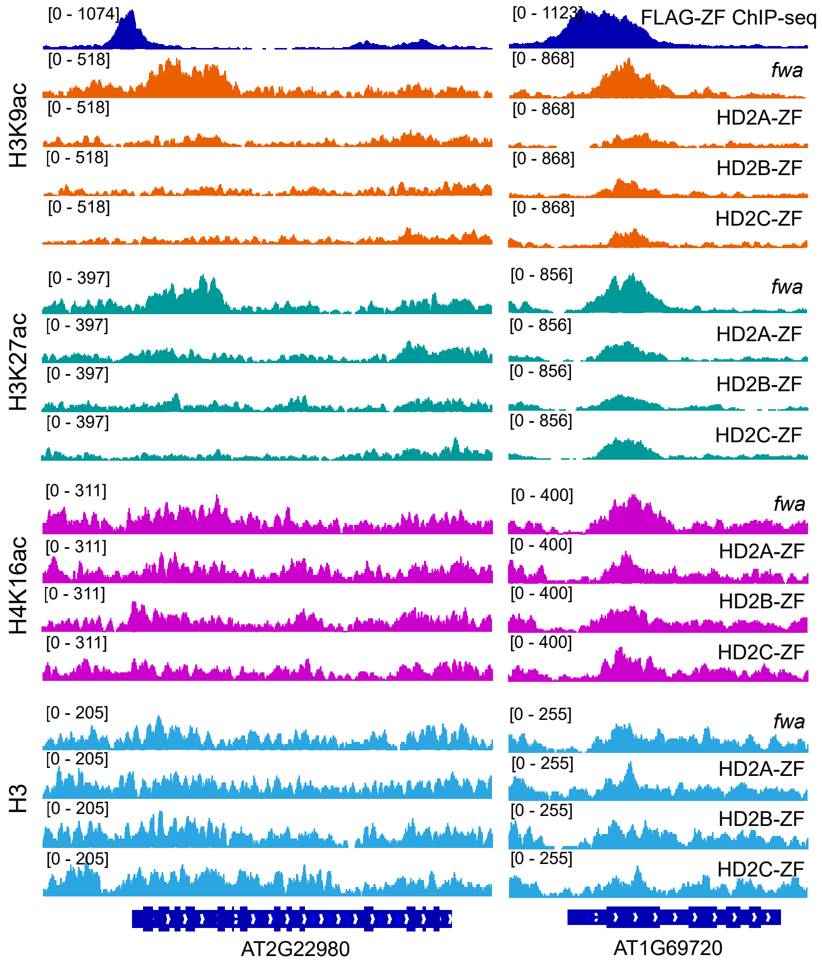


**Supplementary Fig. 4: Target gene silencing by HD2-ZFs and histone H3K9, H3K27, and H4K16 deacetylation.** Screenshots of histone H3K9ac, H3K27ac, H4K16ac, and H3 ChIP-seq signals over two representative ZF off-target sites in *fwa* and T2 transgenic lines of HD2A-ZF, HD2B-ZF, and HD2C-ZF. FLAG-ZF ChIP-seq signals indicate ZF binding sites.

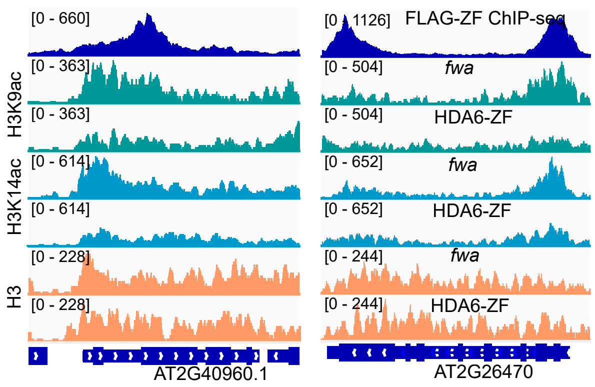


**Supplementary Fig. 5: Target gene silencing by HDA6-ZF and histone H3K9 and H3K14 deacetylation.** Screenshots of histone H3K9ac, H3K14ac, and H3 ChIP-seq signals over two representative ZF off-target sites in *fwa* and HDA6-ZF. FLAG-ZF ChIP-seq signals indicate ZF binding sites.

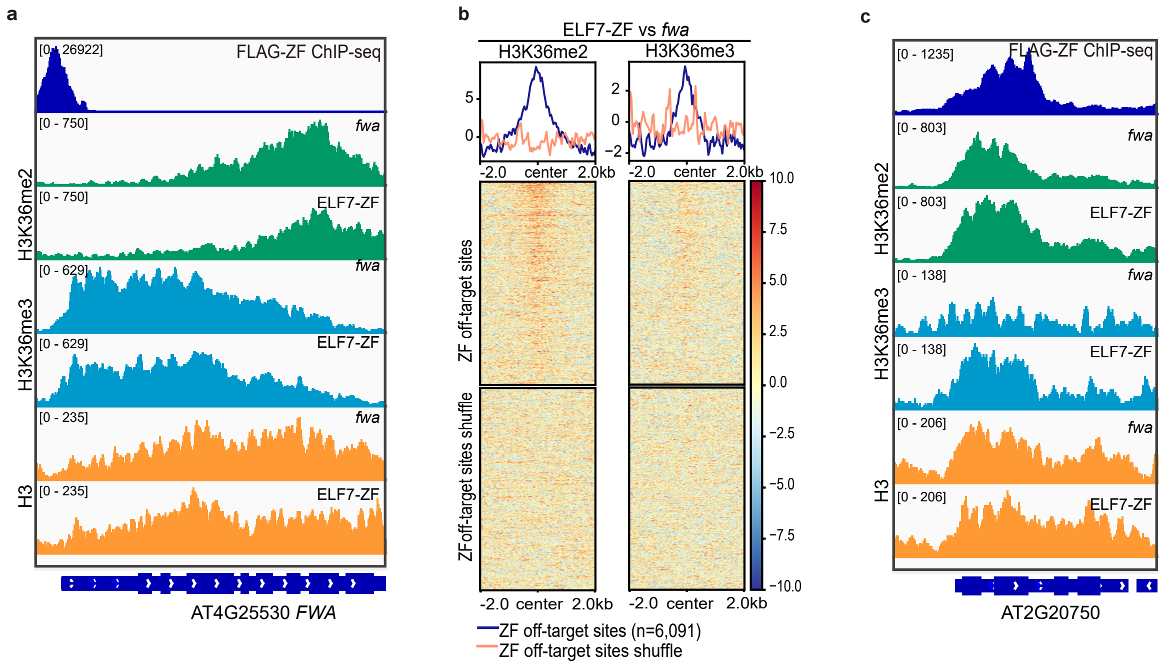


**Supplementary Fig. 6: Target gene silencing by ELF7-ZF cannot be explained by H3K36me2/3 alternations. a,** Screenshots of FLAG-ZF ChIP-seq, and H3K36me2, H3K36me3, and H3 ChIP-seq signals in *fwa* and a representative ELF7-ZF T2 transgenic line over *FWA* region. **b,** Heatmaps and metaplots depicting the normalized H3K36me2 (left) and H3K36me3 (right) ChIP-seq signals in ELF7-ZF versus *fwa* over ZF off target sites (n=6,091). **c,** Screenshots showing the FLAG-ZF ChIP-seq, and H3K36me2, H3K36me3, and H3 ChIP-seq signals in *fwa* and ELF7-ZF over a representative ZF off-target gene.

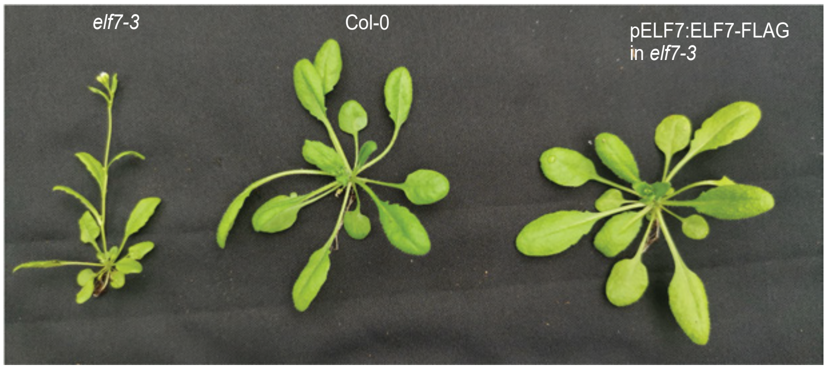


**Supplementary Fig. 7: *elf7-3* mutant was successfully complemented by pELF7:ELF7-FLAG.** Phenotype of 3-4-week-old Arabidopsis *elf7-3* mutant, Col-0 wild type, and pELF7:ELF7-FLAG transgenic line in the background of *elf7-3* mutant.

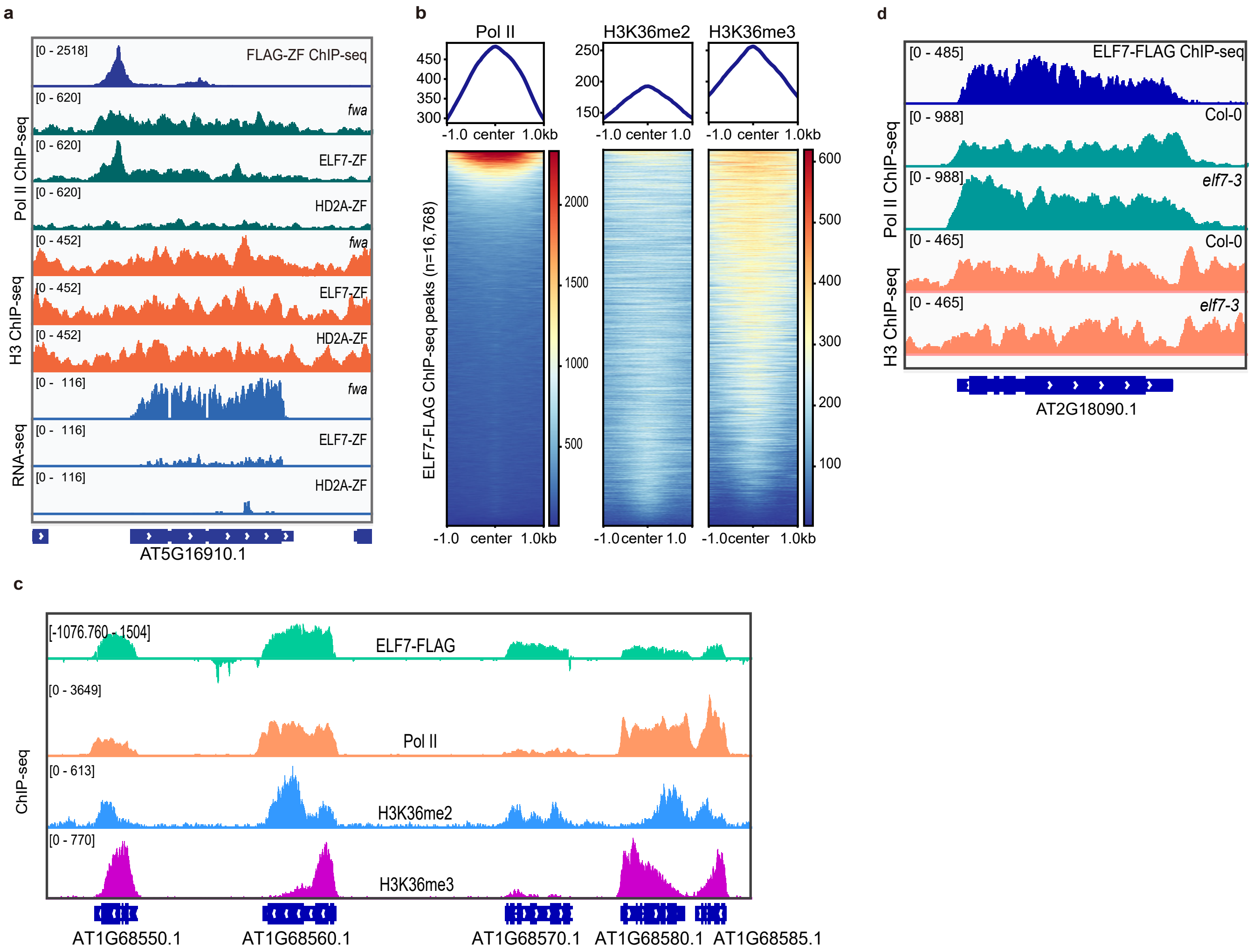


**Supplementary Fig. 8: Target gene silencing by ELF7-ZF. a,** Screenshots of Pol II and H3 ChIP-seq and RNA-seq signals over a representative ZF off-target site in *fwa*, ELF7-ZF, and HD2A-ZF. **b,** Heatmaps and metaplots showing Pol II, H3K36me2, and H3K36me3 ChIP-seq signals over ELF7-FLAG ChIP-seq peaks (n=16,768) in Col-0. **c,** Screenshots of ELF7-FLAG, Pol II, H3K36me2, and H3K36me3 ChIP-seq signals over five representative ELF7 binding genes. **d,** Screenshots of ELF7-FLAG, Pol II, and H3 ChIP-seq signals in Col-0 and *elf7-3* mutants over a representative ELF7 targeting gene.

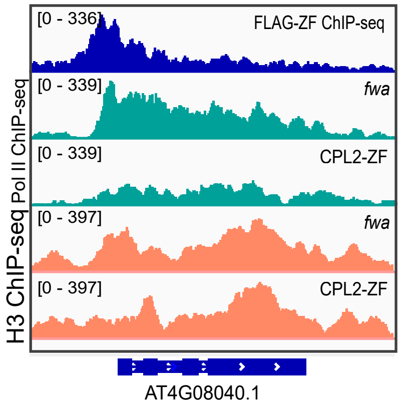


**Supplementary Fig. 9: Target gene silencing by CPL2-ZF and pol II Ser5 dephosphorylation.** Screenshots showing FLAG-ZF and Pol II ChIP-seq signals in *fwa* and T2 transgenic line of CPL2-ZF over a representative ZF off-target site.

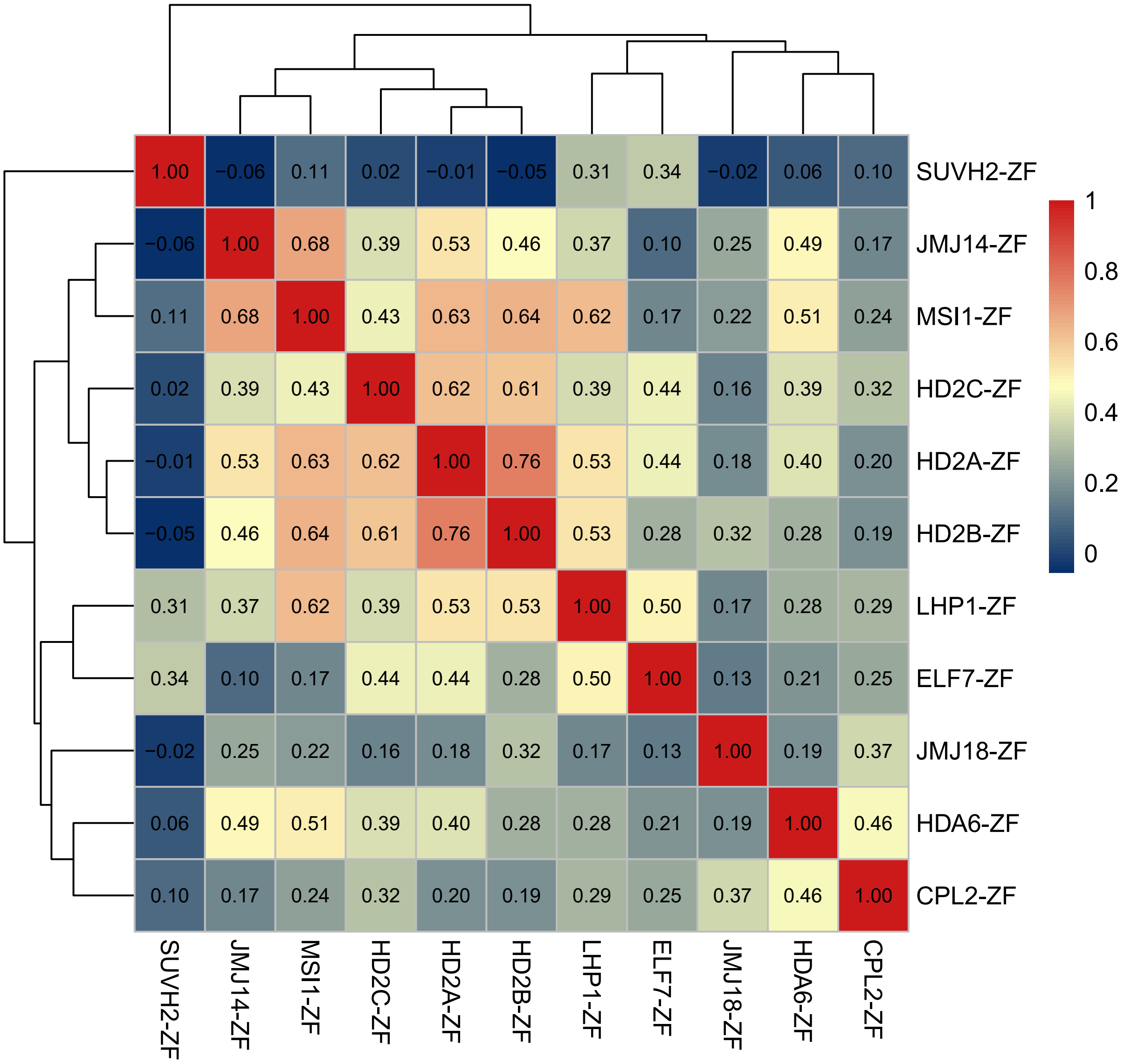


**Supplementary Fig. 10. Overall overlapping of the target genes among all the ZF fusions.** Heatmap displaying Spearman correlation coefficients for all the down regulated ZF target genes (distance to TSS is from -500 to 200 bp, n=805) among all the ZF fusions.

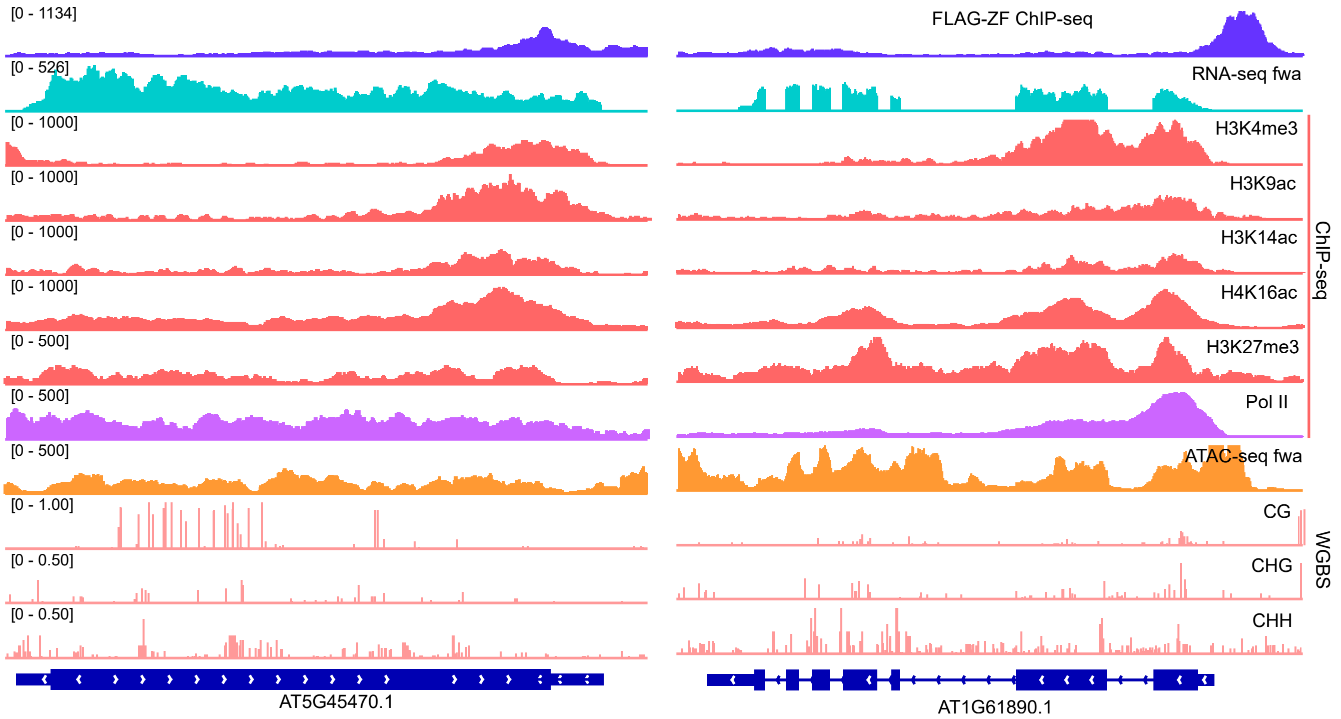


**Supplementary Fig. 11: ZF target genes associated chromatin features are varied.** Screenshots of FLAG-ZF ChIP-seq and RNA-seq in *fwa*, ChIP-seq signals of H3K4me3, H3K9ac, H3K14ac, H4K16ac, H3K27me3, and Pol II in *fwa*, ATAC-seq signals in *fwa*, and CG, CHG, and CHH DNA methylation levels in *fwa* over two representative ZF targeting genes.
